# Aerosolized ApoA1 Nanoparticles Synthesized by Microfluidics Cross the Lung Barrier and Modulate Inflammation

**DOI:** 10.1101/2025.07.09.663869

**Authors:** Marie Laurine Apalama, Luan Givran, Béatrice Navarette, Matthieu Bringart, Pierre Giraud, Philippe Rondeau, Olivier Meilhac, Benoit Allard

**Author notes:** Corresponding authors; these authors contributed equally to this work. DéTROI - Université de la Réunion, 77 Avenue du docteur Jean Marie Dambreville, 97410 Saint Pierre, La Réunion – France.

## Abstract

High-density lipoproteins exert vasculoprotective effects, mainly through apolipoprotein A1, which has led to the development of treatments based on apolipoprotein A1 nanoparticles (A1NPs) administered intravenously, mainly for the treatment of cardiovascular diseases. However, their potential as therapy for lung pathologies has not yet been explored. In this work, we produced A1NPs using microfluidics and characterized their therapeutic potential for lung delivery. Their morphology was characterized by dynamic light scattering and transmission electron microscopy. A1NPs toxicity and cellular uptake were performed on both endothelial (HMEC-1) and epithelial (A549) cells and their anti-inflammatory activity was evaluated on TNF-α-stimulated HMEC-1. A1NPs biodistribution was explored in lung mice after aerosolization and their transcytosis was further investigated using A549 air-liquid interface model. Our results demonstrate that the microfluidic synthesis of A1NPs was reproducible and yielded discoidal particles with sizes ranging from 7-12 nm. A1NPs were internalized by both cells without being cytotoxic and significantly reduced IL-6 expression. Aerosolization resulted in homogeneous distribution in lungs, without causing an immunogenic response. A fraction of A1NPs crossed alveolar epithelial cells both *in vitro* and *in vivo*, paving the way for future therapeutic strategies targeting not only the lungs, but also other peripheral organs. These results are promising for the use of A1NPs as vectors for therapeutic molecules, which could exert synergistic protective effects with Apolipoprotein A1. This is the first study to show the non-invasive administration of A1NPs by aerosolization, which may improve their bioavailability in lungs and appears to be a promising approach for treating lung diseases.

**GRAPHICAL ABSTRACT:** 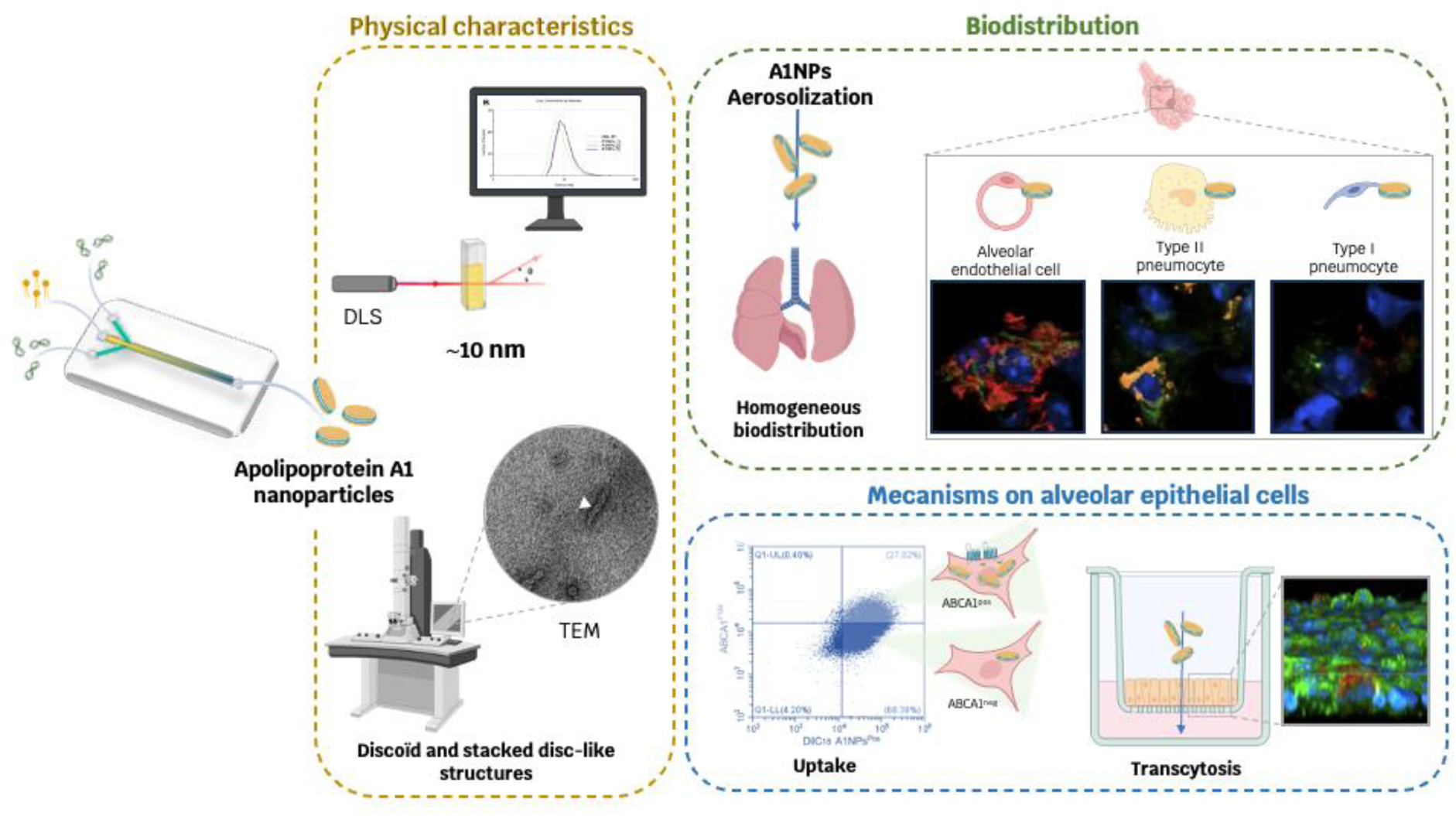

## INTRODUCTION

High-density lipoproteins (HDLs) are complex macromolecules composed of a hydrophobic core with esterified cholesterol and triglycerides, surrounded by phospholipids and proteins. They can adopt a spherical or discoidal conformation depending on their lipid composition^1^. Their high density (1.063 to 1.210 g/mL) is mainly due to their high protein content and their mean size is around 5 to 17 nm according to lipids and protein enrichment^2^. Indeed, HDL particles are composed of several apolipoproteins, among them apolipoprotein A1 (ApoA1) being the major protein^1^. HDLs also contain enzymes such as Lecithin-Cholesterol Acyl Transferase (LCAT) which play a central role in reverse cholesterol transport^3^, giving them anti-atherogenic properties^4^. In addition, HDLs exert anti-inflammatory and antioxidant effects mainly through ApoA1 but also anti-endotoxic effects, linked to their capacity to bind and eliminate lipopolysaccharides, contributing to their overall vasculoprotective action^5–8^.

HDL-cholesterol levels are commonly used as a biomarker of cardiovascular health; a marked decrease being associated with a significant increase in the risk of cardiovascular disease, as initially demonstrated by the Framingham study in 1980^9^. On the strength of their beneficial effects, several pharmaceutical companies have developed HDL-mimetic nanoparticles, known as reconstituted HDL (rHDL) or ApoA1 nanoparticles (A1NPs), since they are composed of ApoA1 and phospholipids. To reduce the cholesterol burden in atherosclerotic plaques, CSL Behring has developed CSL-111 and CSL-112 formulations, composed of phospholipids and ApoA1 isolated from human plasma. These nanoparticles have been evaluated in clinical trials such as ERASE and AEGIS-II, studying the impact of A1NPs on reducing atheroma plaque after intravenous administration^10,11^. Clinical studies testing the effects of reversing atherosclerosis and limiting the recurrence of cardiovascular events have been disappointing^10^. However, A1NPs are currently being evaluated for their anti-inflammatory and endothelioprotective effects in the context of sepsis. We demonstrated that intravenous injection of CSL-111 reduced systemic inflammation, notably through their ability to promote lipopolysaccharide (LPS) clearance in the context of bacterial infection in mice^12^. Our laboratory also confirmed that CER-001 (A1NPs developed by Abionyx, formerly Cerenis) injections in a severe COVID-19 patient decreased circulating inflammatory markers^8^. These anti-inflammatory properties are mainly mediated by ApoA1. Indeed, ApoA1 interacts with the ABCA1 transporter, involved in cholesterol efflux, and leads to the activation of intracellular molecular pathways^14^ leading for example to inhibit the NF-κB signaling pathway in endothelial cells^15^, thereby reducing the production of pro-inflammatory cytokines and chemokines such as IL-6 and MCP-1.

Although A1NPs have been widely studied in the cardiovascular field, no transposition to pulmonary pathologies has yet been established. To target the lung, aerosolization of A1NPs seems far more appropriate than intravenous injection. Apart from inhalation of lipid nanoparticles^16,17^, no one has yet evaluated aerosolization of A1NPs. Various methods for producing these nanoparticles have been developed. The conventional method relies on the use of sodium cholate^18,19^, but its toxicity has led to the development of alternative approaches, notably via microfluidics^20^. This innovative technology enables continuous, rapid production without toxic components, as well as the vectorization of bioactive molecules.

In this study, we produced A1NPs using microfluidics and confirmed their discoidal morphology in a reproducible manner. These nanoparticles were no-cytotoxic to endothelial (HMEC-1) and lung epithelial (A549) cells, and were efficiently internalized. Notably, this uptake was enhanced in the fraction of ABCA1 positive epithelial cells. Their anti-inflammatory property has been confirmed by a significant reduction of IL-6 expression induced by TNF-α stimulation in endothelial cells. *In vivo*, aerosolization of A1NPs led to homogeneous biodistribution throughout the lung, characterized by uptake by type I and II pneumocytes and alveolar endothelial cells. A progressive passage into the systemic circulation from 6 hours post-administration was observed, allowing their biodistribution to peripheral organs, opening up additional therapeutic strategies targeting other pathologies beyond those affecting the lung. Taken together, this study advances our knowledge of the therapeutic potential of A1NPs by opening new perspectives on treatments for respiratory diseases and beyond. The enrichment of A1NPs with therapeutic molecules, particularly hydrophobic, provides insight into new care avenues.

## RESULTS AND DISCUSSION

While therapies based on apolipoprotein A1 nanoparticles (A1NPs) have failed to achieve the expected effects in cardiovascular diseases, their therapeutic potential in lung diseases has not been explored. Here, we have combined the fields of physics and biology to shown that anti-inflammatory A1NPs can be homogeneously aerosolized in the lungs, paving the way for new therapeutic strategies for lung diseases.

### Production and physical characterization of A1NPs

A1NPs were reconstituted using a single-step, self-assembly method in a single layer, 3-inlet microfluidic device (Figure 1A). As previously shown by Kim et al., this technique allows the production of reproducible and homogeneous batches of A1NPs^20^. While Kim et al. performed their nanoparticle synthesis in phosphate buffer saline, we opted for reconstitution directly in a buffer designed to preserve A1NPs by limiting their oxidation (TEN buffer). It is also worth noting that we used a five-fold higher concentration of ApoA1, enabling us to obtain nanoparticles of the expected majority size without the need for additional purification steps. Dynamic light scattering (DLS) analysis confirmed the reproducibility of these productions. These results indicate that the average size of A1NPs was around 10 nm, similar to that of plasma HDLs^1^ (Figure 1B). The structural organization of A1NPs was observed under a transmission electron microscope (Figure 1C). This observation revealed that A1NPs formed stacked disc-like structures (discoidal shape), also known as “rolls”, similar to those observed for microfluidically synthetized Apo A1 nanoparticles of Kim et al.^20^ and already described for plasmatic preβ-HDL^21^. The overall morphology of A1NPs was comparable to that of preβ-HDL demonstrating the ability of our laboratory to generate biological nanoparticles, as previously done by pharmaceutical groups. CSL-111 and CSL-112 are nanoparticles made from human ApoA1 and soy-derived phospholipids and have been evaluated in clinical trials^22^, making them a benchmark in terms of morphological and size characteristics. Yet, in the context of coronary artery disease, these nanoparticles did not produce the expected atheromatous plaque reduction effect, underlining the need for optimizations to improve their therapeutic efficacy^22^. One proposed solution is to enrich these nanoparticles with bioactive molecules. For instance, Moreno et al. showed that high-density lipoproteins (HDLs) enriched with alpha-1-antitrypsin, significantly reduced neutrophil elastase-induced pulmonary emphysema in mice, compared with native, plasma isolated HDLs^23^. Thus, the use of A1NPs as vectors for therapeutic molecules represents a promising approach requiring further investigation to optimize their therapeutic potential. Microfluidic offers significant advantages for this type of enrichment, whether of proteins, synthetic molecules or lipids^24^. Controlled flow rates and channel dimensions in the micrometer range promote molecule assembly, facilitating the incorporation of therapeutic compounds into nanoparticles, without using additional chemicals.

**Figure 1.**
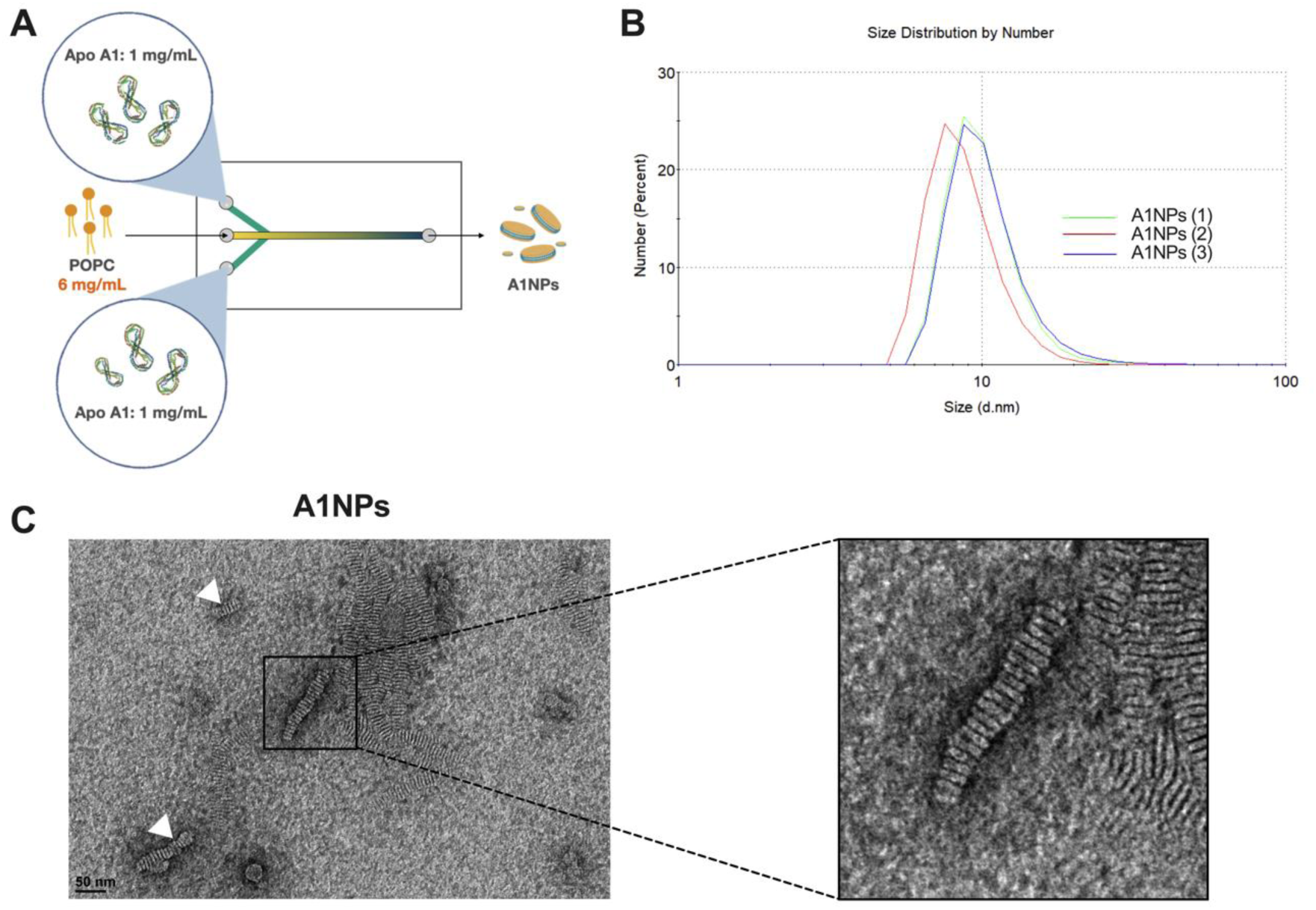
Physical characterization of Apolipoprotein A1 nanoparticles. (A) A1NPs were produced by microfluidic using a chip with 2 inlets for Apolipoprotein A1 (ApoA1) injection at 0.8 mL/min and 1 inlet for phospholipids (POPC) injection at 0.1 mL/min. (B) Three independent productions of A1NPs were characterized by dynamic light scattering to determine their size (7-12 nm). (C) Electron transmission microscopy was used to determine the shape of A1NPs. White triangles indicate stacked disc-like structures, also known as “rolls”.

### A1NPs are internalized by cells without cytotoxicity

To assess the safety profile of A1NPs, cell viability tests were conducted across a range of A1NP concentrations. HMEC-1 and A549 cells were incubated with A1NPs at concentrations from 0.015 mg/mL to 1 mg/mL for 24 hours, followed by an MTT assay (Figure 2A,B). The results indicate no significant difference between stimulated and unstimulated cells, suggesting A1NPs have no impact on the viability of either cell type. To visualize A1NPs internalization into cells, confocal microscopy was performed on both HMEC-1 and A549 cells using anti-ApoA1 antibodies. Observations show that after 6 hours, cells have internalized the A1NPs, with an increase in labeling intensity observed after 24 hours (Figure 2C). In addition to MTT assay, these findings indicate that the nanoparticles do not alter cell morphology and integrity. It should be emphasized that nanoparticles not only bind to the cell surface, but are also internalized. Indeed, Silver et al. demonstrated that hepatocytes incubated with HDLs at 4°C only led to binding. At 37°C, results indicate an active uptake process^25,26^. Given that the uptake of HDLs is mediated by the scavenger receptor class B type I (SR-BI) and the ATP binding cassette subfamily A member 1 (ABCA1)^27^, one may argue that A1NPs internalization is also dependent on these receptors. Alveolar epithelial cells (type I and type II) as well as A549 cells do express ABCA1^28,29^ but whether A1NP-mediated uptake is similar to that of endothelial cells^27,30^ is unclear. To clarify the potential role of ABCA1 in A1NP uptake by A549 cells, we incubated cells with fluorescent A1NPs (DilC_18_ staining) for 6 hours and analyzed cells by flow cytometry. We observed that about 95% of A549 cells were positive for DilC_18_-A1NP. However, only approximately 27% of A549 cells were positive for ABCA1, but this population captured a higher amount of DilC_18_-A1NP, as evidenced by a significant increase in the mean fluorescence intensity of DilC_18_ compared to ABCA1-negative A549 cells (Figure 2D,E and S1). This result confirms that a fraction of A549 cells do express ABCA1 in unstimulated condition and that ABCA1 is, at least in part, involved in A1NP uptake. Other receptors, also expressed by A549 cells, such as SRB-1^31^, could also be involved in A1NPs uptake, thereby explaining that 95% of cells are able to internalize these nanoparticles. Interestingly, the dual role of ABCA1 in repressing inflammation while maintaining cholesterol homeostasis represents a promising therapeutic target for inflammatory lung diseases in the future^29^.

**Figure 2.**
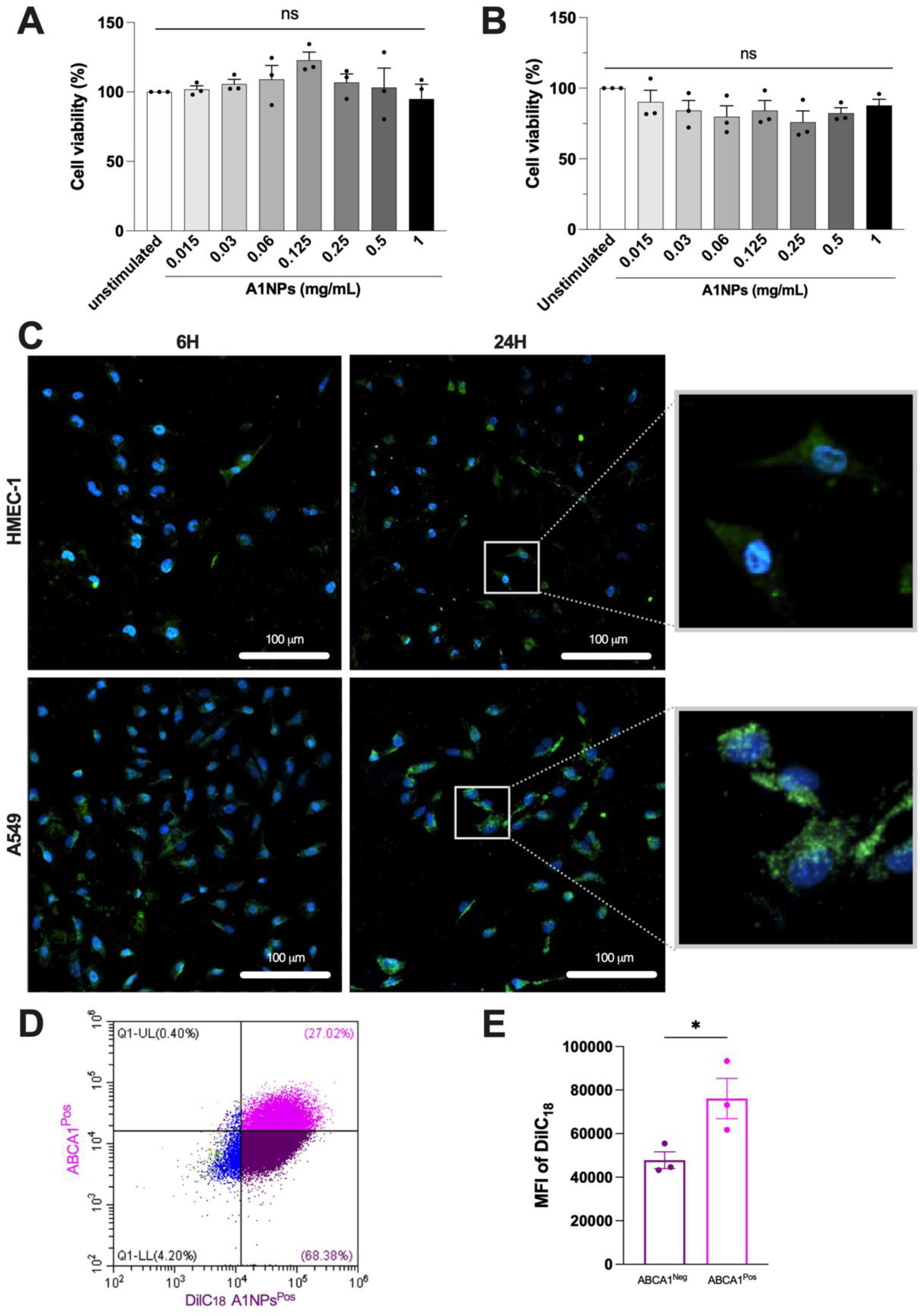
Apolipoprotein A1 nanoparticle cytotoxicity and uptake assay. (A,B) MTT assay on endothelial (HMEC-1, A) and alveolar epithelial cell lines (A549, B), 24 hours after A1NP incubation. (n=3-5 independent experiments; ANOVA; Tukey’s multiple comparison test; * p<0.05, ** p<0.01, **** p<0,0001). (C) A1NPs [0.05mg/ml] are taken up by both cell lines after 6 and 24 hours of incubation. A rabbit anti human ApoA1 antibody was used for labelling A1NPs (green) and cell nuclei were stained with DAPI (blue). (D,E) Uptake of A1NPs by ABCA1-positive A549 cells. (D) A dot plot illustration for double staining with DilC_18_-A1NPs and ABCA1 and (E) the mean fluorescence intensity of DilC_18_ in ABCA1 negative and ABCA1 positive cells. (n=3 independent experiments; Paired t-test; * p < 0.05).

### A1NPs display anti-inflammatory properties

Endothelial cells were stimulated with 2.5 ng/mL of TNF-α to induce an inflammatory response and co-stimulated with A1NPs. Following 6 hours of incubation, we assessed the gene expression of the pro-inflammatory mediator IL-6 (Figure 3A). After 16 hours, IL-6 protein levels in the culture medium were quantified by ELISA (Figure 3B). Both IL-6 mRNA and protein levels increased about 3-fold under inflammatory conditions compared to untreated controls. However, treatment with A1NPs significantly reduced IL-6 expression at both the transcriptional and protein levels, indicating an anti-inflammatory effect. These findings demonstrate that A1NPs are biologically functional and exhibit anti-inflammatory activity in HMEC-1 cells under TNF-α-induced inflammatory conditions. This assay on endothelial cells is a classical hallmark to appreciate the anti-inflammatory properties of HDLs and mimetics^32^. It has been shown that ApoA1 binding to ABCA1 may trigger the expression of tristetraprolin, which subsequently promotes the degradation of inflammatory cytokine mRNA in response to LPS, including IL-6, through its 3’-UTR AREs^33^.

**Figure 3.**
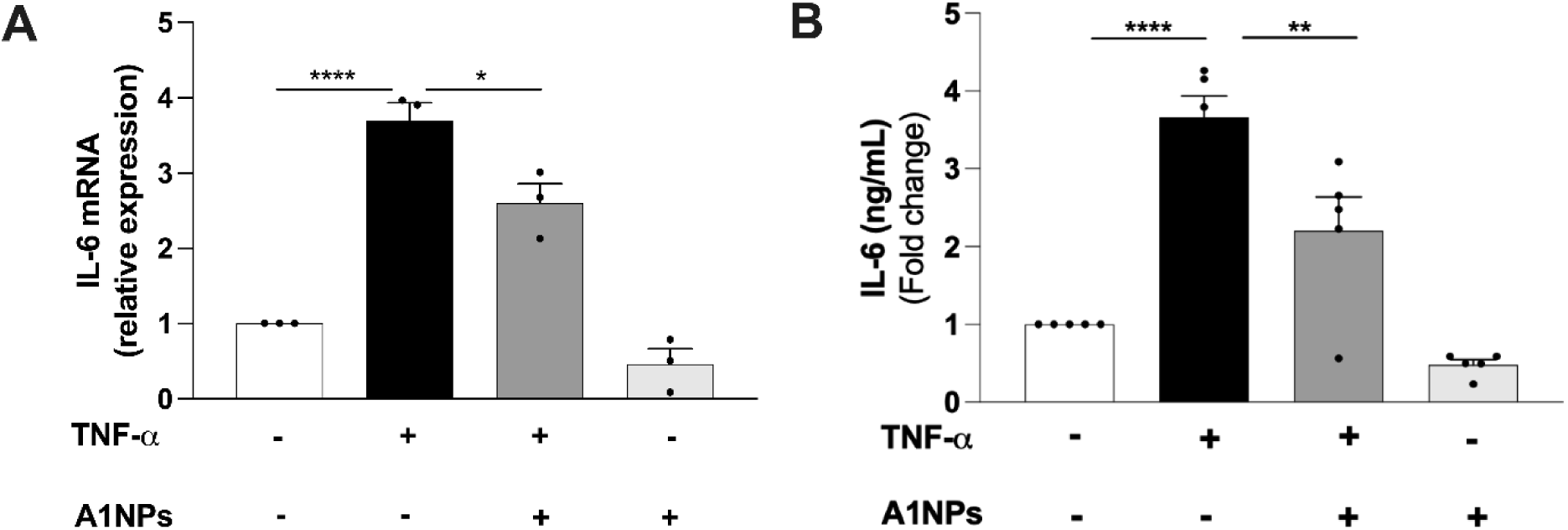
Anti-inflammatory properties of A1NPs. A1NPs [0.05 mg/ml] significantly decrease TNF-α-induced IL-6 at mRNA (A) and protein levels (B) in HMEC-1 cells. (n=3-5 independent experiments; ANOVA; Tukey’s multiple comparison test; * p<0.05, ** p<0.01, **** p<0,0001).

### A1NPs are homogeneously distributed in the lung after aerosolization

To investigate the biodistribution of A1NPs following their administration by aerosolization, A1NPs were first fluorescently labelled with DilC_18_ (Figure 4A). Since the lung is closely linked to the capillary network, we also quantified the passage of A1NPs in the bloodstream. A1NPs reached a peak in plasma 6 hours after administration, before decreasing 12 hours later, but their detection persisted after 24 hours (Figure 4B). At 6 hours post-administration, homogeneous red fluorescence was detected throughout the lung parenchyma of mice that had received DilC_18_-A1NPs while no fluorescence was observed in control mice aerosolized with PBS (Figure 4C). This uniform biodistribution of A1NPs persisted at 24 hours, which is also consistent with A1NP plasma kinetics (Figure 4B). Interestingly, red fluorescence was also detected in the liver and kidneys at 24 hours (Figure S2). These observations suggest that A1NPs-DilC_18_ behave similarly to HDL particles, with elimination via hepatic and renal pathways. No significant variation in the body weight of the mice was observed throughout the experiment, supporting the absence of *in vivo* nanoparticle toxicity (Figure S3A). Moreover, additional experiments have demonstrated the absence of immunogenicity in mice given A1NPs on days 0, 1 and 12 (Figure S3B). Surprisingly, other organs, such as the brain and the spleen, were also enriched in A1NPs after their passage into the bloodstream (Figure S4), opening up therapeutic prospects targeting these organs. Although the lung remains the main organ targeted by aerosolization, this non-invasive route could also be considered for the treatment of pathologies characterized by chronic inflammation in peripheral organs.

**Figure 4.**
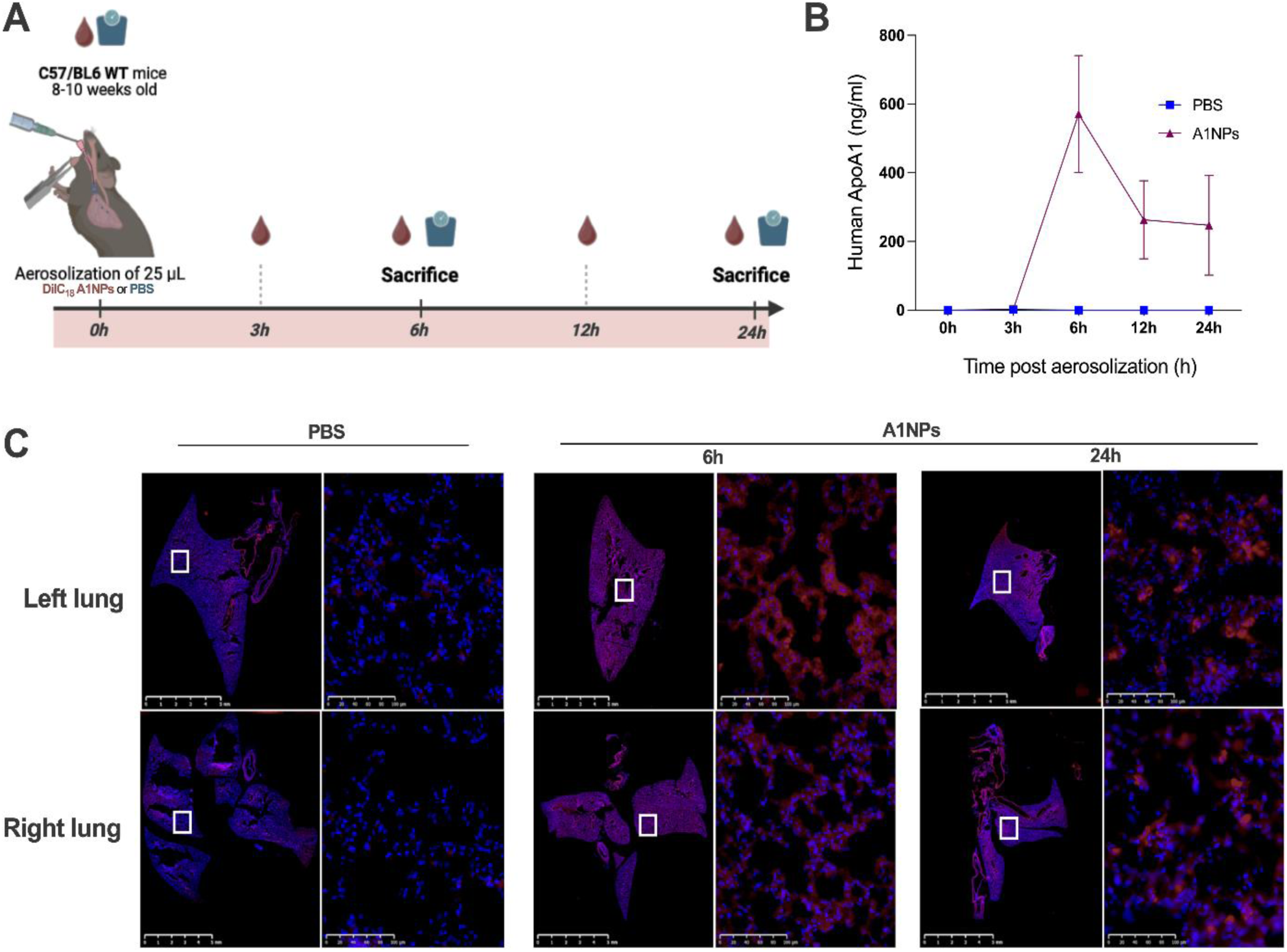
Biodistribution of A1NPs after aerosolization. (A) Experimental design for studying the biodistribution of A1NPs administered intratracheally by aerosolization. (B) Determination of human ApoA1 concentration in mouse plasma at different time points after aerosolization: 0h, 3h, 6h, 12h and 24h (n=6 for PBS, n=10 for A1NPs). (C) A1NPs labeled with DilC18 dye (red) reach the left and right lungs 6 hours after aerosolization and persist in both lungs for up to 24 hours. Representative illustration of 6 mice treated with PBS and 10 mice with A1NPs-DilC18.

To further evaluate the precise localization of A1NPs, we performed co-labeling between ApoA1 and different cell types specific to lung tissue. We observed co-localization of ApoA1 with cell-specific markers (AGER: type I pneumocytes; SFPTC: type II pneumocytes; EMCN: vascular endothelial cells) (Figures 5 and S5). These different structural lung cell types were able to internalize A1NPs, opening up interesting therapeutic perspectives. These include intracellular application of A1NPs potentially enriched with therapeutic molecules that act intracellularly, such as siRNAs. This A1NP’s broad spectrum of pulmonary penetration makes it the vector of choice for lung diseases.

**Figure 5.**
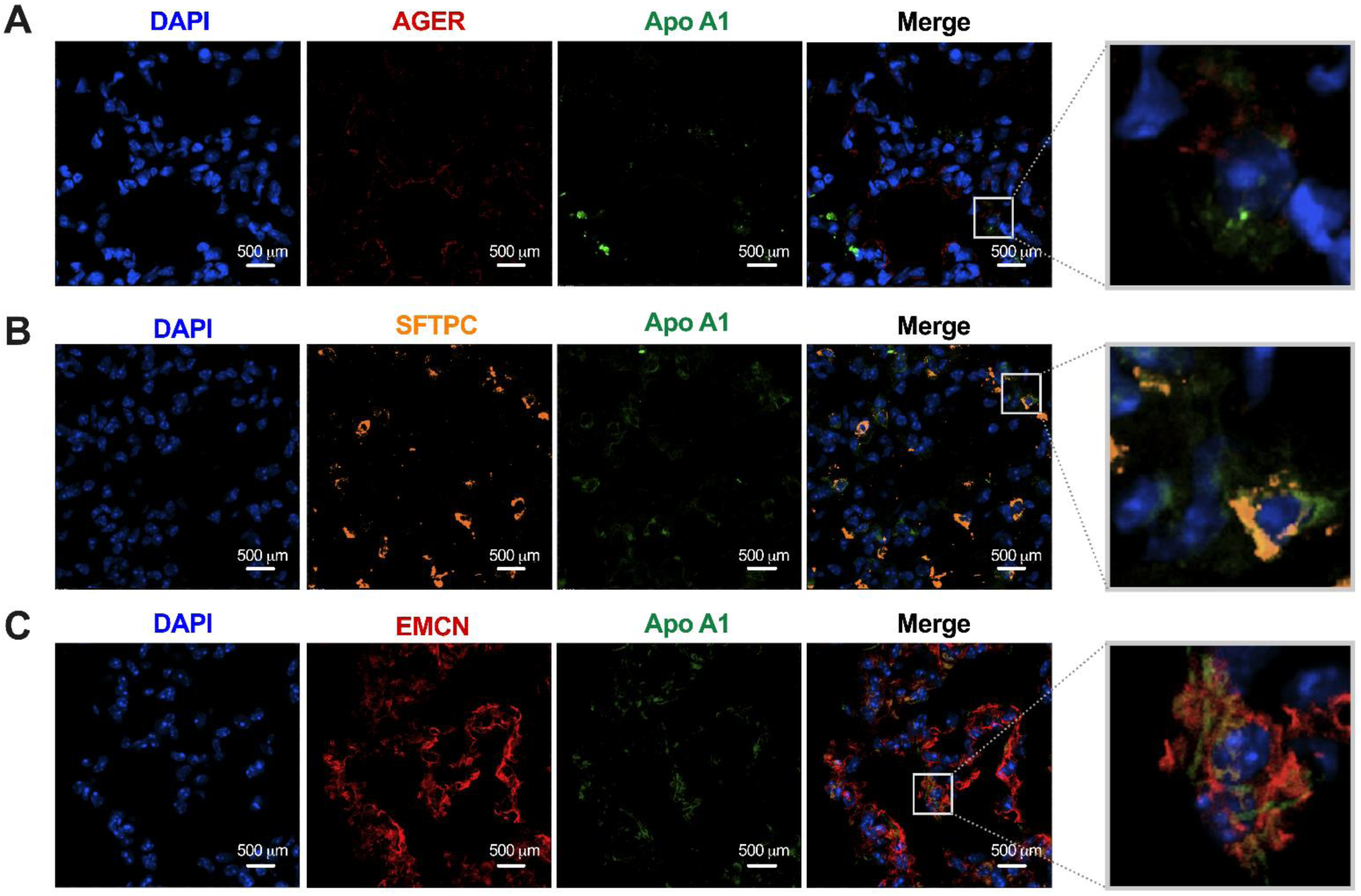
Cellular localization of A1NPs after aerosolization. Immunofluorescence of lung sections from mice 6 hours after A1NP administration. ApoA1 appears in green and cell nuclei are stained with DAPI (blue). (A) AGER (red), specific to type I pneumocytes. (B) SFTPC (orange), marker for type II pneumocytes. (C) EMCN (red), characteristic of pulmonary endothelial cells. Representative illustration of 10 mice with A1NPs-DilC18.

### A1NPs pass through an epithelium grown at an air-liquid interface

Previous results *in vivo* suggest a progressive transfer of A1NPs into the bloodstream. The nanoparticles may cross the alveolar barrier in the lung, being initially internalized by alveolar epithelial cells before reaching endothelial cells, allowing access to the vascular compartment. To further explore the passage of A1NPs through an epithelium, we set up an air-liquid interface (ALI) model of lung epithelial cells using inserts (Figure 6A). According to previous characterization of ALI culture of A549 alveolar epithelial cells, this model reconstitutes epithelial layers with the expression of markers of both alveolar epithelial type I and type II cells^34^. After apical addition of A1NPs, the inserts were incubated for 6 hours. A progressive passage of A1NPs was observed over time in these experimental conditions (Figure 6B). In another set of experiments, we made sure that after the assay, the permeability of both A1NPs and without A1NPs epithelium was the same, ruling out the possibility that stimulation may alter this parameter (Figure S6). Confocal microscopy confirmed the presence of ApoA1 in the cytoplasm of epithelial cells (Figure 6C,D and S7). A deeper understanding of the mechanisms involved in the transepithelial passage of A1NPs would be relevant, to determine whether this is a process of transcytosis or other alternative mechanisms. ABCA1 is known to facilitate the interaction and internalization of pre-β HDL particles. This mechanism has been extensively characterized in endothelial cells^30^. Given that A1NPs exhibit structural and functional similarities to pre-β HDL, one may argue that their cellular internalization is also mediated by ABCA1. Our flow cytometry results on ABCA1 expression showing an increase uptake in ABCA1 positive cells support a model in which ABCA1 partially mediates A1NP internalization, although additional receptors may contribute to uptake through other specific mechanisms. Consistent with existing literature, discoidal ApoA1 particles have been shown to preferentially interact with ABCA1 to facilitate lipid acquisition^30^. In contrast, the SR-B1 receptor is known to recognize lipid-rich spherical HDL particles. The ABCG1 transporter also plays a role in lipid efflux and may be implicated in HDL trafficking^15^. It is interesting to note that Moreno et al. observed that intravenous injection of HDL in mice with elastase-induced emphysema led to increased HDL recruitment in the lungs compared to control mice^23^. This study suggests that A1NP may be preferentially recruited to inflamed tissues. Collectively, our data suggest that the microfluidically produced A1NPs mimic the biological behavior of circulating discoidal HDL particles and may use similar uptake pathways.

**Figure 6.**
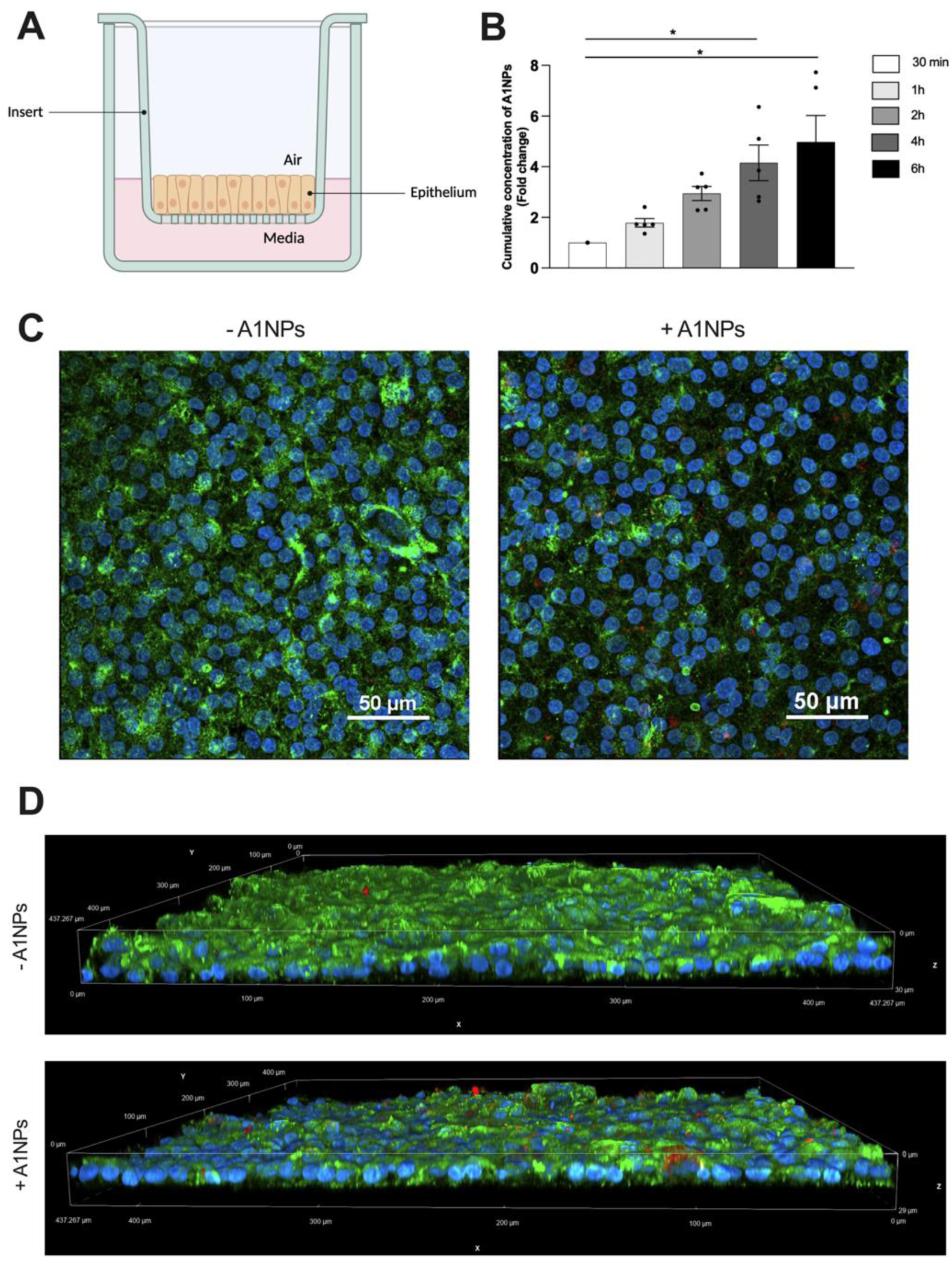
Passage of A1NPs through A549 epithelial cells grown in air-liquid interface. (A) Schema showing the experimental setup used to study A1NPs migration through lung epithelial cells cultured in air/liquid interface (ALI). (B) Quantification of A1NPs transcytosis under ALI conditions, measured by ELISA specific for human ApoA1, in the basolateral medium. The cumulative concentration of A1NPs in the basolateral medium is determined at different incubation times: 30 min, 1h, 2h, 4h and 6h. (C) Immunofluorescence analysis of ALI membrane after 6h of transcytosis with 0.5 mg/mL A1NPs. Phalloidin (green) labels the actin cytoskeleton, cell nuclei are stained blue, and ApoA1 appear in red. (D) Three-dimensional visualization of the ALI membrane under different experimental conditions.

## CONCLUSION

In this study, we report the successful development of apolipoprotein A1-based nanoparticles (A1NPs) produced by microfluidics and designed for pulmonary delivery. Our results demonstrate that A1NPs possess favorable physicochemical characteristics, including a reproducible discoidal morphology and nanoscale dimensions (7–12 nm), comparable to clinically evaluated HDL mimetics. *In vitro*, A1NPs were efficiently internalized by both endothelial and alveolar epithelial cells, with uptake enhanced in ABCA1-positive populations, and displayed no cytotoxicity across a range of concentrations. Importantly, A1NPs retained biological functionality, as evidenced by their significant anti-inflammatory effects on TNF-α-stimulated endothelial cells, with a reduction in IL-6 expression at both transcript and protein levels.

Upon aerosolization in mice, A1NPs exhibited homogeneous pulmonary biodistribution, reaching both lobes and persisting in lung tissue for up to 24 hours without triggering detectable immunogenicity. Furthermore, we confirmed their transcytosis across alveolar epithelial barriers both *in vitro* and *in vivo*, with a progressive passage through the epithelium. This dual capacity for local action and systemic translocation highlights the versatility of A1NPs not only as lung-targeted agents but also as systemic drug delivery vehicles following non-invasive administration.

Together, these findings position A1NPs as a promising nanoplatform for the treatment of respiratory diseases, with the potential to be enriched with bioactive molecules for synergistic therapeutic effects. Future investigations will focus on optimizing A1NPs for targeted delivery of anti-inflammatory or antioxidant agents, and evaluating their efficacy in relevant models of acute and chronic lung inflammation. This work opens new perspectives for the translation of HDL-mimetic nanoparticles into pulmonary medicine.

## MATERIALS & METHODS

### A1NPs production and characterization

A1NPs are synthesized from plasma apolipoprotein A1 (ApoA1) (plasma was obtained from the French blood national agency, EFS-LR agreement number #2018001378) and commercial phospholipids (2-oleoyl-1-palmitoyl-sn-glycero-3-phosphocholine; 42773-500MG, Sigma). The production is carried out on a microfluidic chip with three input channels. The central inlet is for phospholipids prepared at a concentration of 6 mg/mL in ethanol at an injection flow rate of 0.1 mL/min (Darwin Microfluidics, SeryngeONE, Connection Kit 01). The other two inlets are used to inject ApoA1 at a concentration of 1 mg/mL in TEN buffer (10 mM TRIS, 1mM EDTA, 150 mM NaCl) at an injection rate of 0.8 mL/min. The solution obtained at the microfluidic chip outlet is centrifuged at 12,000 g for 15 minutes to sediment the aggregates. A1NPs are then concentrated and washed with a TEN buffer using a 10 kDa cut-off concentrator (Corning).

### Dynamic light scattering characterization

A1NPs were characterized by dynamic light scattering (DLS) spectroscopy as described previously^35^.

### Electron microscopy characterization

Transmission electron microscopy analyses were conducted at the Center for Quantitative Imaging Lyon East (CIQLE, University of Lyon 1, Lyon, France). The morphological characteristics and size of the nanoparticles were assessed using negatively stained samples, imaged with a Gatan Orius 600 CCD camera (Gatan, USA) on a LaB6 JEOL JEM-1400 transmission electron microscope (JEOL, Japan) operating at an accelerating voltage of 120 kV. For sample preparation, 300-mesh copper grids coated with a carbon film (Delta Microscopies, France) were glow-discharged for 30 seconds (Balzers SCD 040, Liechtenstein) to render the carbon surface hydrophilic, facilitating sample adhesion. Nanoparticle suspensions were subsequently deposited onto the treated grids, followed by negative staining using a 2% aqueous solution of uranyl acetate. After complete air drying, the grids were mounted onto a single-tilt holder and introduced into a JEOL JEM-2100 transmission electron microscope (JEOL, Japan), equipped with a cryo pole piece and operated at 120 kV. Image acquisition was performed using a Gatan SC600A CCD camera (Gatan, USA).

### Cell Culture and stimulation

HMEC-1 cells, a human microvascular endothelial cell line (ATCC# CRL-3243), were cultured in MCDB 131 medium (P04-80057, Pan Biotech) supplemented with 10% fetal bovine serum (FBS) (ST30-3302, Pan Biotech), 100 units/mL Penicillin-100µg/mL Streptomycin (P06-07100, Pan Biotech), 250µg/mL Amphotericin-B (P06-01100, Pan Biotech), 10 ng/mL Epidermal Growth Factor (EGF) (E9644; Sigma-Aldrich, USA), 1 µg/mL Hydrocortisone (Sigma), and 10 mM L-Glutamine (Pan Biotech). A549 cells, a human alveolar epithelial cell line (ATCC# CCL-185), were routinely cultured in RPMI 1640 medium (P04-22100, Pan Biotech) supplemented with 10% heat-inactivated FBS (ST30-3302, Pan Biotech) and 100 units/mL Penicillin-100µg/mL Streptomycin (P06-07100, Pan Biotech). Both cell lines were maintained in a humidified incubator at 37°C with 5% CO₂ and subcultured when they reached 90% confluence. During stimulation with A1NPs, the respective media were supplemented with 10% delipidated FBS for HMEC-1 and 5% delipidated FBS for A549.

### A1NP Cytotoxicity

The cytotoxicity of A1NPs on each cell line was evaluated using the 3-(4,5-dimethyl-thiazol-2-yl)-2,5-diphenyl tetrazolium bromide (MTT) assay. 96-well plate were seeded with 10,000 cells/well. After 24hours of incubation, the medium was removed and the cells were stimulated with 100 µL of A1NPs at concentrations ranging from 1 mg/mL to 0.015 mg/mL for 24 hours. 10µL of MTT reagent (M2128-5G, Sigma-Aldrich) diluted in PBS was added to each well for five hours, resulting in a final MTT concentration of 0.5 mg/mL. The plate was then centrifuged at 500g for 4 minutes at 25°C. The medium was removed and replaced with 200 µL of dimethyl sulfoxide (DMSO) (UD8050-B, Euromedex) to dissolve the formazan crystals. Absorbance was measured at 560 nm using a CLARIOStar Plus plate reader (BMG Labtech).

### A1NP uptake by endothelial and epithelial cells

8-well Labtek chamber slides were seeded with 50,000 cells/well for each cell line studied. Once cell confluence reached 80%, the wells were washed with PBS solution and were serum-deprived for 3 hours. Cells were stimulated with A1NPs at a concentration of 0.05 mg/mL, or with their respective media for 6 hours and 24 hours. Cells were washed three times with PBS, then fixed with 4% paraformaldehyde (PFA) for 15 minutes. Slides were washed once with PBS for 5 minutes, then twice with PBS containing 0.05% Triton for 10 minutes. Cells were then blocked for 1 hour in PBS 0.05% triton 2% BSA. Primary anti-human ApoA1 antibody (Calbiochem) was diluted 1:500 in PBS 0.05% triton 0.2% BSA and incubated with the cells overnight at 4°C. After 3× 10-minute washes, the cells were incubated with DAPI (1μg/mL, Sigma) mixed with an Alexa 488 goat anti-rabbit secondary antibody at 1:1000 dilution for 1 hour at room temperature. After 5x-10-minute washes, the slides were mounted with fluorescent medium and images were captured with a confocal microscope (Nikon Eclipse Ti2).

### Flow Cytometry

A total of 200,000 A549 epithelial cells were seeded in 6-well plates. After 24 hours, the culture medium was replaced with a lipid-depleted medium for an additional 24-hour period. Subsequently, cells were either stimulated or not with DilC_18_ A1NPs at a concentration of 0.05 mg/mL for 4 hours, under lipid-free conditions. Following stimulation, cells were detached using Accutase (25-058-CI, Corning) and immediately placed on ice. Cell suspension was centrifuged at 300g for 5 minutes. The supernatant containing Accutase was discarded, and the resulting cell pellet was resuspended in FACS buffer (PBS supplemented with 0.1% BSA). After a second centrifugation, the supernatant was removed, and Fc receptor blocking reagent (anti-CD16/CD32; 553141, BD Bioscience) was added at a concentration of 1 µg per 10⁶ cells. The suspension was incubated for 10 minutes at 4°C. Cells were then washed with FACS buffer and incubated with the human ABCA1 Alexa Fluor 488 conjugated antibody (NB100-2068G, NovusBio) at 1 µg per 10⁶ cells, for 30 minutes in the dark at 4°C. Following a final wash with PBS, the cells were resuspended and analyzed by flow cytometry (Cytoflex, Beckman Coulter).

### A1NP anti-inflammatory properties

The anti-inflammatory activity of A1NPs was assessed in the TNF-α-stimulated HMEC-1 cell line by RT-qPCR and ELISA.

### RT-qPCR

50,000 cells were seeded in a 24-well plate. When the cells reached 70% confluence, they were washed and serum-deprived (0% FBS) for 3 hours. Cells were stimulated with A1NPs at a concentration of 0.05 mg/mL for 6 hours, with or without addition of TNF-α (2.5 ng/mL). After stimulation, cells were lysed and RNA was extracted with RNeasy Plus Mini kit (74136, Qiagen) and quantified by Nanodrop (BMG Labtech). Reverse transcription was carried out using the NxGen M-MuLV reverse transcriptase (30222-1, Lucigen) according to the manufacturer’s standard protocol. Quantitative PCR was performed using the Blastaq Green 2x qPCR MasterMix (G892, Abm). The transcription levels of IL-6 were measured using the following primers: forward 5’-ACCCCCAGGAGAAGATTCCA-3’, reverse 5’-GCCTCTTTGCTGCTTTCACA-3’. The data were normalized against GAPDH (forward 5’-AGCCACATCGCTCAGACAC-3’, reverse 5’-GCCCAATACGACCAAATCC-3’) and RNA polymerase II (RNApol2) (forward 5’-CGAGAAGGTCTCATTGACACAG-3’, reverse 5’-ACCACCTGGTTGATGGAGTTCC-3’).

### ELISA

70,000 cells/wells were seeded in a 12-well plate. When the cells reached 70% confluence, they were washed and serum-deprived (0% FBS) for 3 hours. Next, cells were stimulated with A1NPs at a concentration of 0.05 mg/mL for 16 hours, with or without the addition of TNF-α (2.5 ng/mL). Supernatant was collected for IL-6 quantification using a human IL-6 ELISA Ready-SET-Go assay (Thermofisher). Absorbance was measured using a CLARIOstar plate reader (BMG Labtech) at 450-570 nm.

### Aerosolization in mice

C57BL/6 mice (8-10 weeks; male female ratio 1:1) were fed ad libitum with standard laboratory chow and water. All animal experiments were approved by the local ethics committee and Ministry of Higher Education and Research (APAFIS #53302-2025011413372952 v4). To visualize A1NPs in the lung, the nanoparticles were first incubated with 200 μg of DilC18 at 37°C overnight with agitation at 250 RPM. Free DilC18 was removed by ultracentrifugation. The density of the A1NPs-DilC18 was adjusted to 1.23 g/mL, and the particles were then recovered by a first layer consisting in KBr at a density of 1.21 g/mL and a second layer at 1.063 g/mL. Following ultracentrifugation at 252,000g for 18 hours at 4°C, the A1NPs-DilC18 were collected. ApoA1 concentration in A1NPs was determined by BCA protein assay (Sigma). A1NPs-DilC18 were prepared at 8mg/mL. For intratracheal aerosolization of A1NPs, mice were anesthetized with isoflurane (2.5%), followed by an intraperitoneal (i.p.) injection of ketamine (90 mg/kg) and xylazine (4.5 mg/kg) to deepen anesthesia. They were positioned on a panel inclined at 45° for intratracheal instillation, with a light source to visualize the tracheal orifice. Using a microsprayer aerosolizer, model YAN 30012 (Yuyan Instruments), a volume of 25 µL was administered between the vocal cords. Mice received either PBS solution or A1NPs-DilC18 (8 mg/ml). Blood was collected before and after aerosolization at 3, 6, 12 and 24 hours. 4 mice (1 PBS and 3 A1NPs-DilC18) were sacrificed at 12 hours and 4 mice (1 PBS and 3 A1NPs-DilC18) were sacrificed at 24 hours. After transcardiac perfusion with PBS and then 4% PFA, the lungs were then fixed for 24 hours in 4% PFA, kept in 30% sucrose solution overnight, and then frozen in OCT at −80°C.

### Immunostaining

Frozen sections (10 µm) were obtained using a cryostat (Leica CM1520; Leica Biosystems). OCT was eliminated with PBS and tissue section were incubated with DAPI (1 μg/mL) at RT for 20 minutes. Ibidi mounting medium was used to see fluorescence and images were obtained unsing a Nanozoomer S60 digital slide scanner (Hamamatsu).

To assess the cell types capable of internalizing A1NPs, co-labeling involving ApoA1 and specific markers was performed. Tissue sections were first subjected to antigen unmasking in sodium citrate (pH 6), held at 80°C for 30 minutes. Once cooled to room temperature, the slides were washed with PBS, then blocked for 90 minutes in PBS 0.1% Triton - 2% BSA. Then, slides were incubated for a further 30 minutes in PBS, 0.1% Triton - 0.2% BSA with Fc Block at 0.025 mg/mL. Slides were then incubated overnight at 4°C with specific primary antibodies. For ApoA1 detection, a mouse anti-human ApoA1 primary antibody diluted 1:200 was used. In parallel, specific antibodies were applied: a mouse anti-AGER rat antibody diluted 1:100 for type I pneumocytes (MAB 1179-500; R&D Systems), a mouse anti-SFTPC rabbit antibody diluted 1:200 for type II pneumocytes (10774-1AP; Proteintech), and a mouse anti-EMCN goat antibody diluted 1:100 for endothelial cells (AF4666-SP; R&D Systems). After five minutes washes with PBS 0.1% Triton, the slides were incubated with secondary antibodies coupled to suitable fluorochromes, including a donkey anti-mouse Alexa Fluor 488, a goat anti-rat Alexa Fluor 594, goat anti-rabbit Alexa Fluor 647, and donkey anti-goat Alexa Fluor 594, all diluted at 1:1000. After this step, the slides were washed three times with PBS 0.1% Triton, then incubated with DAPI at 1 µg/mL for 15 minutes for nuclei staining.

Finally, after 3 washes in PBS 0.1% Triton, slides were mounted with IBIDI medium. Observations and acquisitions were made using a confocal microscope (Nikon Eclipse Ti2).

### Air/liquid interface model

A total of 500,000 A549 cells were seeded into polycarbonate cell culture inserts with pore-size of 0.4 μm (PIHP01250; Millipore) pre-coated with 70 µg/mL of type I rat tail collagen (354236, Corning, USA). After 24 hours, the medium in the apical compartment was removed and the medium at the basolateral side was replaced 3 times per week for 2 weeks^34^.

### Transcytosis

To assess the potential of A1NPs to cross a reconstituted epithelial barrier, the inserts were exposed to 0.05 mg/mL of A1NPs. Basolateral media were collected after 30 minutes, 1 hour, 2 hours, 4 hours, and 6 hours and analyzed with a human ApoA1 ELISA assay (3710-1HP-2, Mabtech). Absorbance was measured using a CLARIOstar plate reader (BMG Labtech) at 450-570 nm. In another set of experiments, A1NPs-exposed inserts were washed with PBS (top and bottom), then fixed with 4% paraformaldehyde (PFA) for 15 minutes on the top and the bottom. Inserts were washed twice with PBS and could either be stored at 4°C for 1 week or used directly. Membranes of insert were cut and washed with PBS 0.5% Triton for 5 minutes. Cells were blocked with PBS triton 0.5% BSA 4% for 30 minutes. Primary anti-human ApoA1 antibody (178422; Calbiochem) was diluted 1:500 in PBS triton 0.5% BSA 1% and incubated for 45 minutes at RT. After 2 washes with PBS, membranes were incubated with Alexa 594 goat anti-rabbit secondary antibody at 1:1000 dilution for 45 minutes at RT. After 2 washes with PBS, membranes were incubated with Alexa Fluor 488 phalloidin at dilution 1:2000 and DAPI (1μg/mL) for 45 minutes at RT. Membranes were mounted with fluorescent medium and images were captured with a confocal microscope (Nikon Eclipse Ti2).

### Permeability assay

Following 6 hours of transcytosis, the inserts were retrieved and transferred into 300 µL of complete culture medium without phenol red (P04-16516, Pan Biotech). Subsequently, 100 µL of dextran labeled with fluorescein isothiocyanate (FITC-dextran) 70 kDa (Sigma) at a concentration of 1 mg/mL was applied to the apical compartment. The inserts were incubated at 37 °C for 40 minutes. Post-incubation, the basolateral medium was collected for each experimental condition. The fluorescence intensity was quantified using a spectrophotometer with excitation/emission wavelengths set at 490 ± 15 nm and 530 ± 30 nm (BMG Labtech). The concentration of FITC-dextran (70 kDa) in the basolateral compartment was determined using a standard calibration curve ranging from 0.25 mg/mL to 0.008 mg/mL. Result was expressed as the relative concentration, calculated as the ratio between the initial concentration applied at T₀ and the concentration measured at T₄₀ minutes (Ct_40_/Ct_0_).

### Statistics

All statistical tests were performed on Graphpad Prism 5 software (Graphpad Software, San Diego, CA). Results were displayed as mean ± SEM values of repeated independent experiments. Statistical tests used were ordinary one-way ANOVA with Tukey’s multiple comparisons test or paired T test. Results were considered statistically significant when p<0.05.

