## Supplemental data for "Aerosolized ApoA1 Nanoparticles Synthesized by Microfluidics Cross the Lung Barrier and Modulate Inflammation"

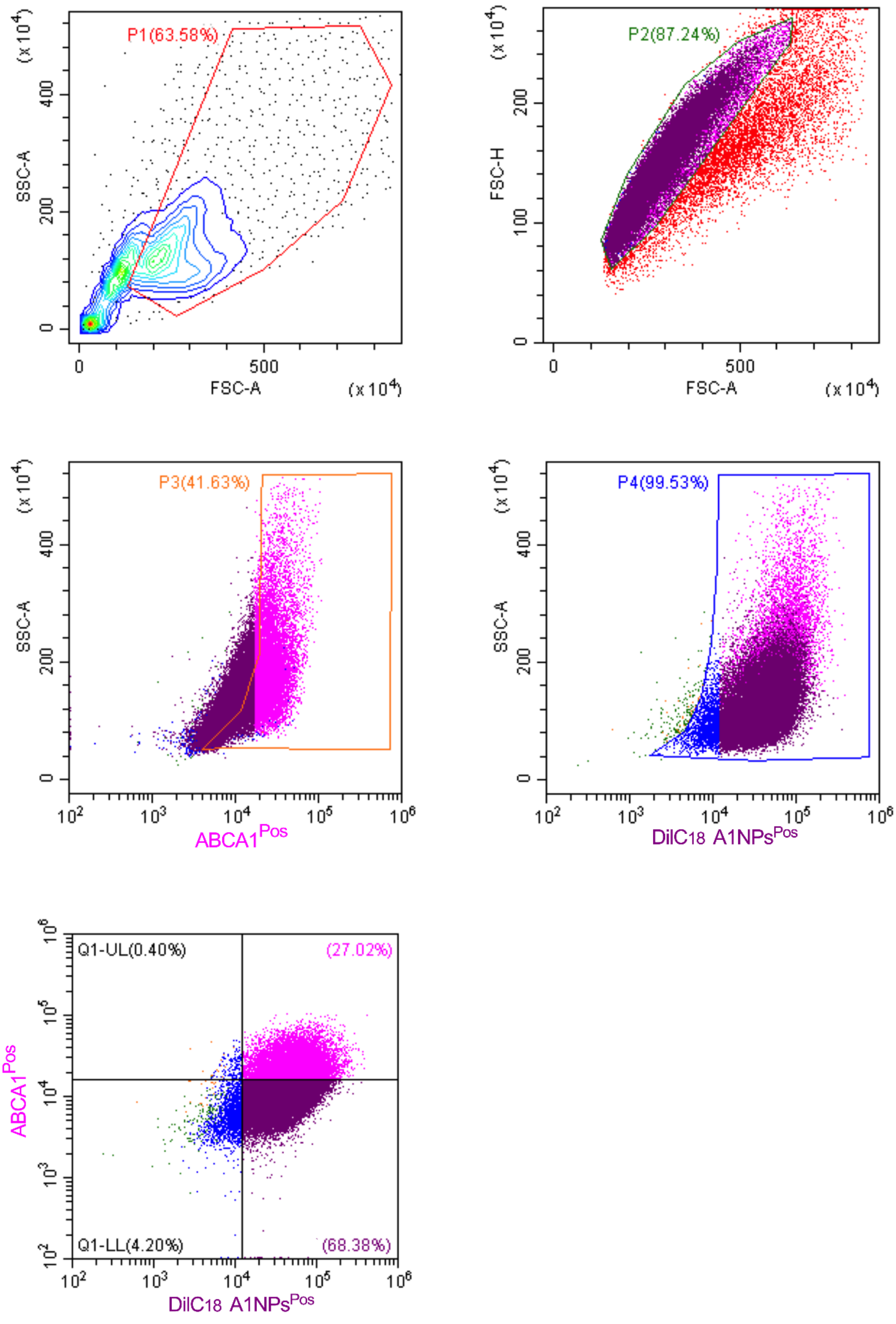


**Figure S1. Gating strategy used to identify ABCA1 positive cells in the A549 cell population. Debris were first excluded based on forward and side scatter parameters (P1), followed by the elimination of doublets cells (P2). ABCA1 positive cells were identified in gate P3, and DilC18-positive cells were gated in P4. Double staining for ABCA1 and DilC18 is shown in the last dot plot. Flow cytometry data were acquired using the CytoFLEX system and analyzed with CytExpert software.**

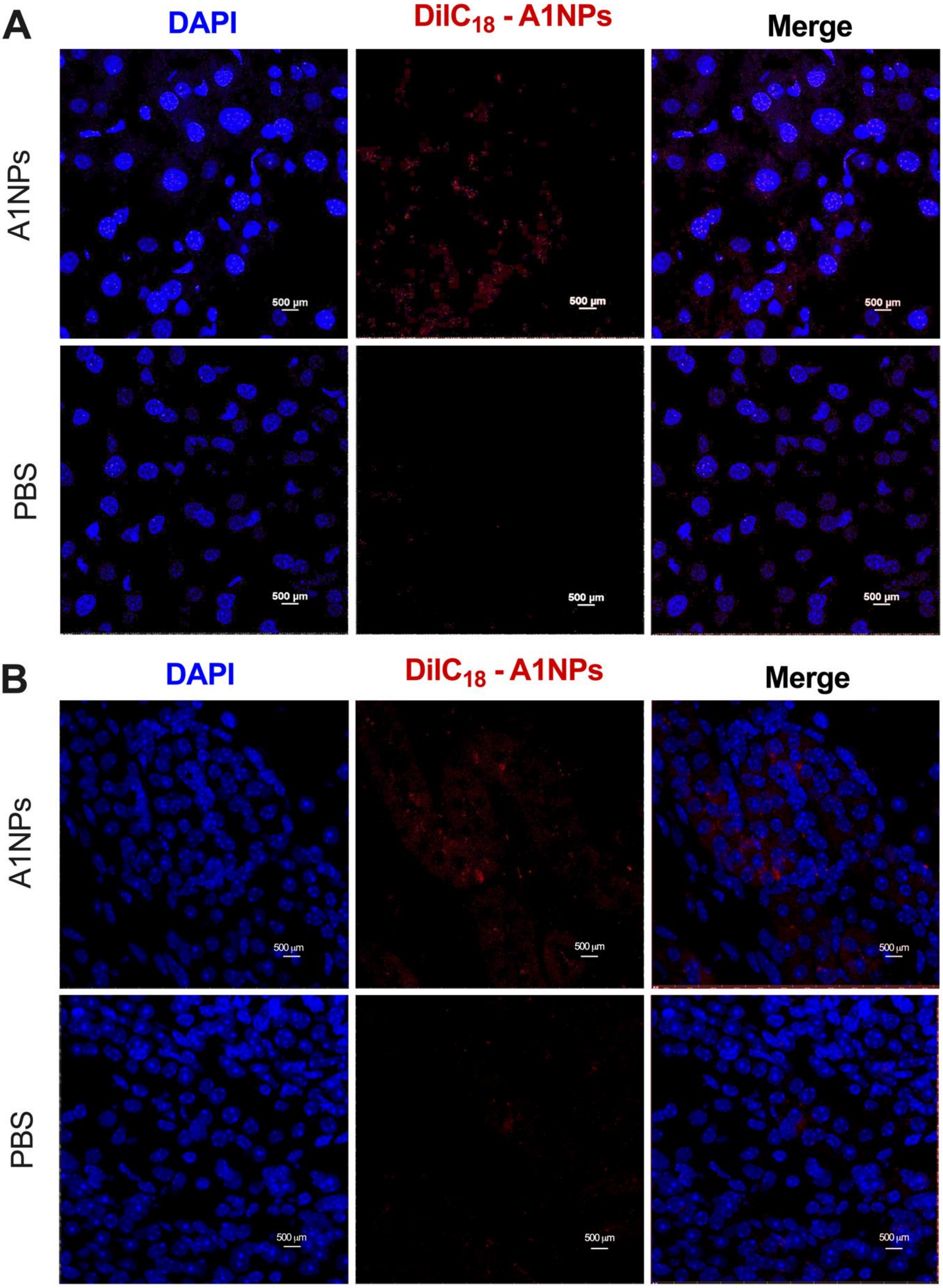


**Figure S2. A1NP biodistribution in detoxifying organs. A1NPs labeled with DilC18 dye (red) reach liver (A) and kidneys (B) 24h hours after aerosolization. Representative illustration of 6 mice treated with PBS and 10 mice with A1NPs-DilC18.**

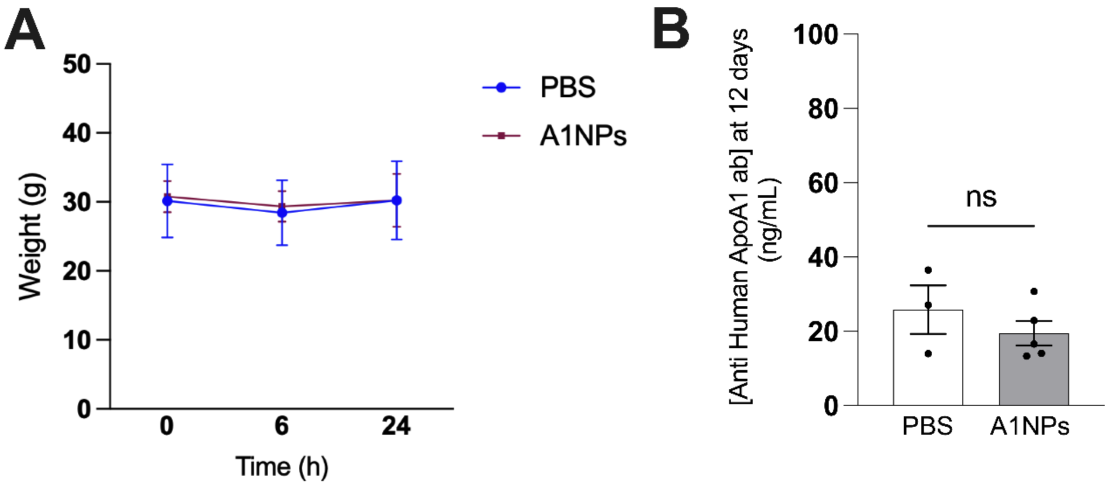


**Figure S3. Changes in mouse weight and production of anti-ApoA1 antibodies.** **(A) Curve showing change in mouse weight over time. (B) Concentration of anti-human ApoA1 antibodies measured in mice 12 days after the second administration of A1NPs.**

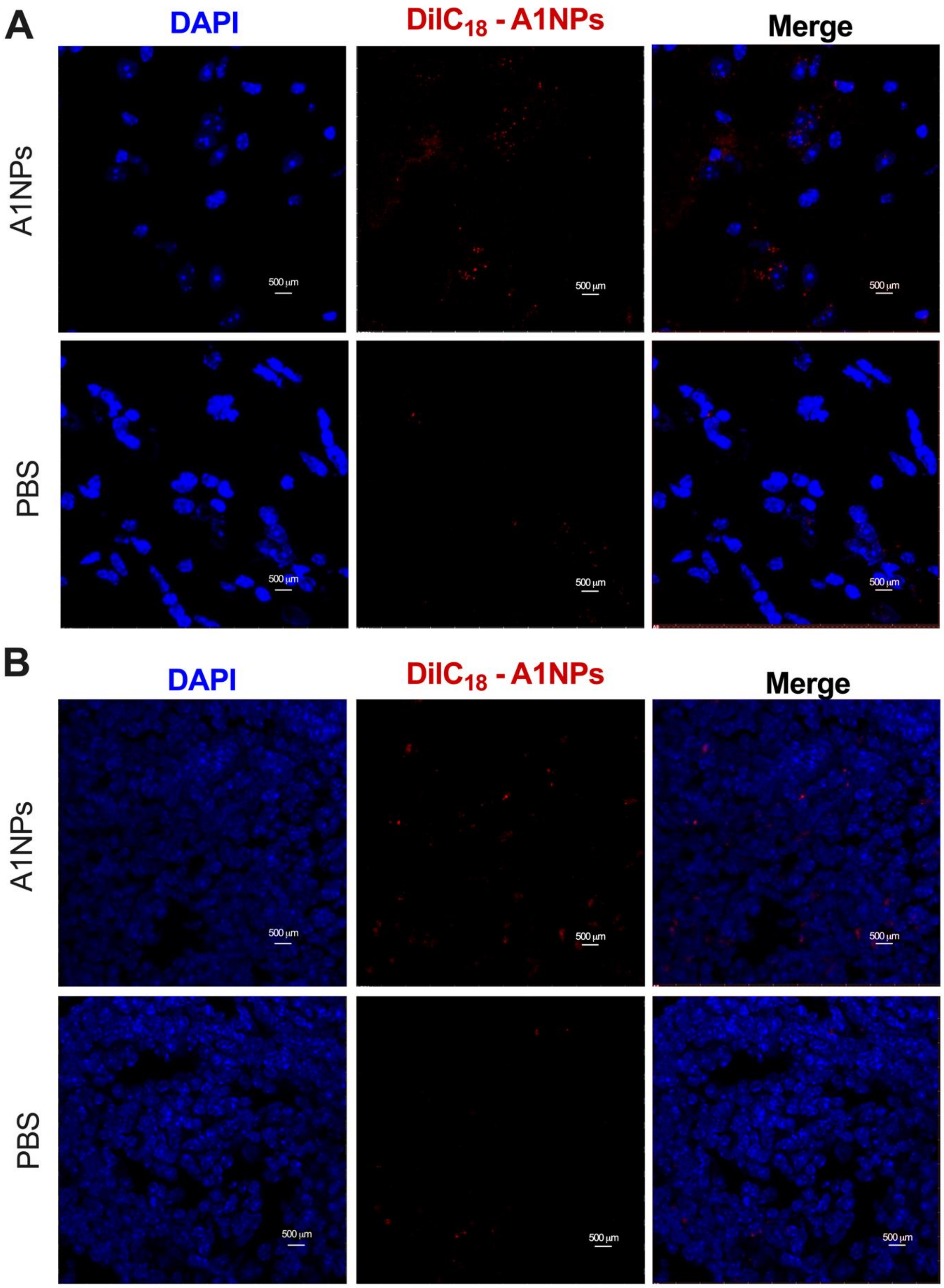


**Figure S4. A1NP biodistribution in other peripheral organs. A1NPs labeled with DilC$_{18}$ dye (red) reach the brain (A) and the spleen (B) 24 hours after aerosolization. Representative illustration of 6 mice treated with PBS and 10 mice with DilC$_{18}$ A1NPs.**

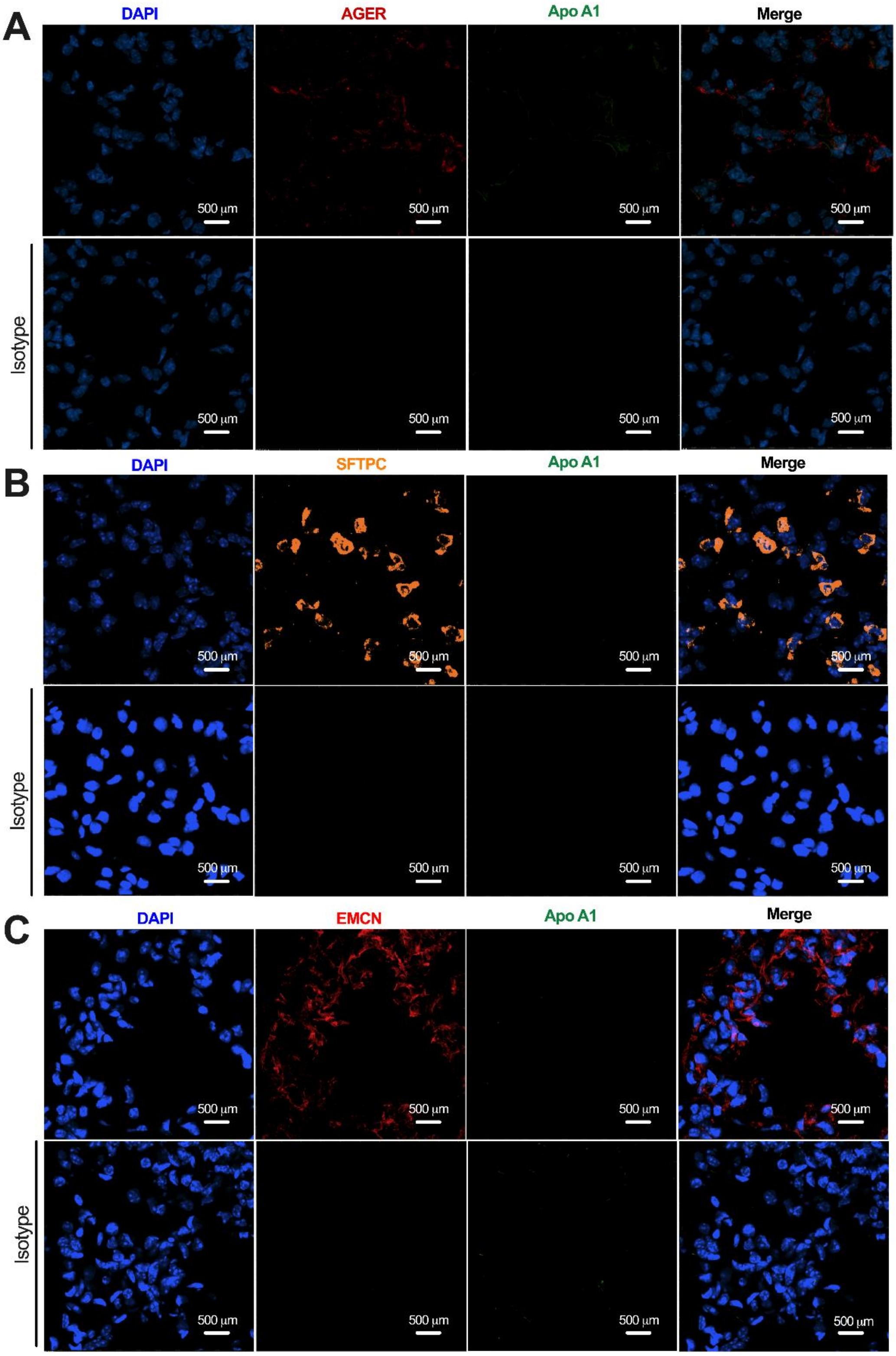


**Figure S5. Control panel of immunofluorescence of lung sections from mice 6 hours after A1NP administration. ApoA1 appears in green and cell nuclei are stained with DAPI (blue). (A) AGER (red), specific to type 1 pneumocytes. (B) SFTPC (orange), marker for type 2 pneumocytes. (C) EMCN (red), characteristic of pulmonary endothelial cells. First line is PBS-treated mice with primary antibody and second line is PBS-treated mice with isotype staining.**

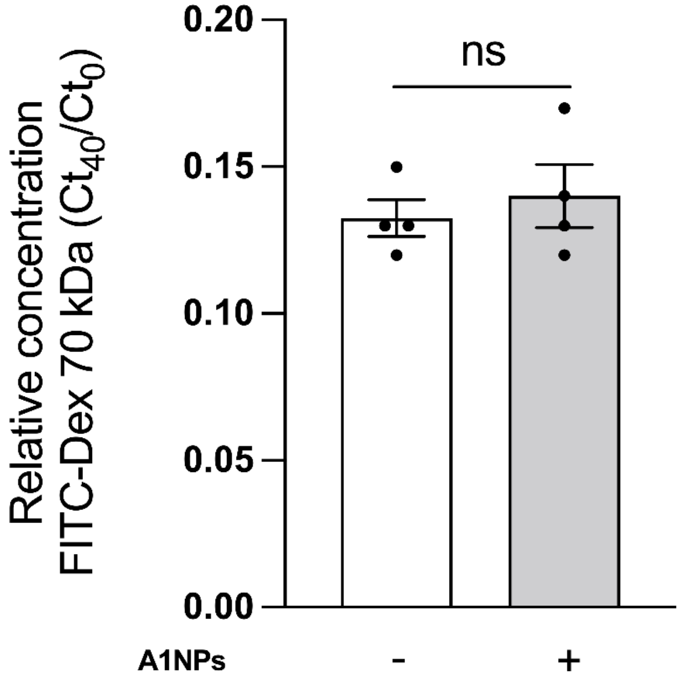


**Figure S6. Passage of FITC-Dextran-70 kDa through the epithelium. Relative concentration of FITC-Dextran-70 kDa passing through the epithelium after 6 hours exposure to A1NPs in apical. $C_{t40}$ corresponds to the concentration of FITC-Dextran-70 kDa found in the basolateral medium after 40 minutes of incubation, while $C_{t0}$ represents the initial concentration of FITC-Dextran-70 kDa placed apically.**

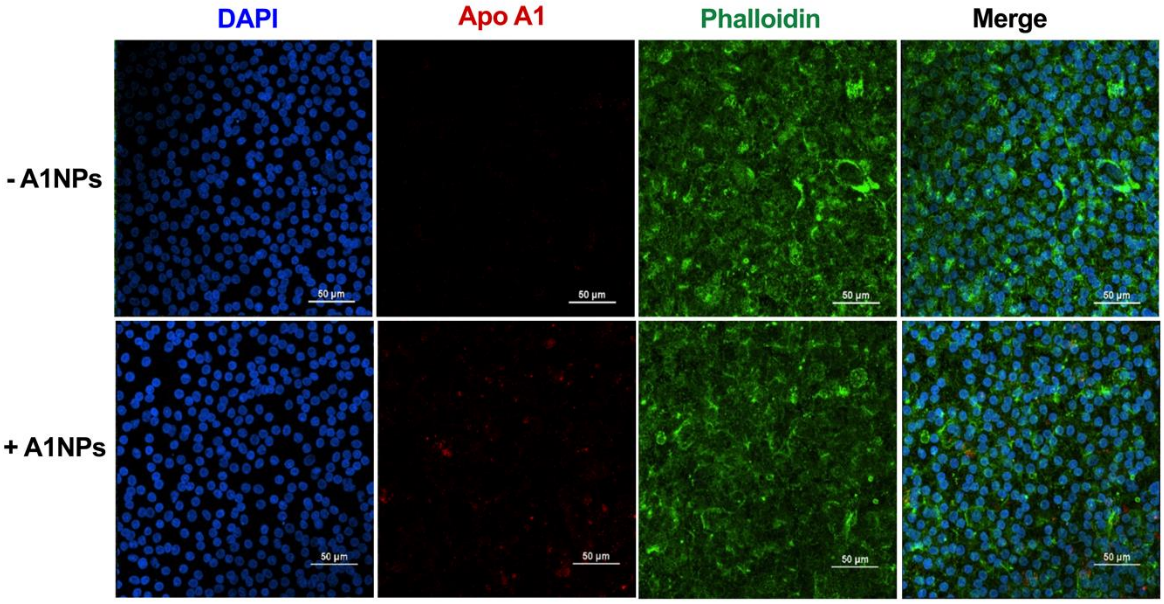


**Figure S7. Immunostaining of A549 in air-liquid interface after A1NP transcytosis. Immunofluorescence analysis of ALI membrane after 6 hours of transcytosis with 0.5 mg/mL A1NPs. Phalloidin (green) marks the actin cytoskeleton, cell nuclei are stained blue, and ApoA1 appear in red.**
